# Accumulation of mutations in genes associated with sexual reproduction contributed to the domestication of a vegetatively propagated staple crop, enset

**DOI:** 10.1101/2020.04.01.020750

**Authors:** Kiflu Gebramicael Tesfamicael, Endale Gebre, Timothy J March, Beata Sznajder, Diane E. Mather, Carlos Marcelino Rodriguez Lopez

**Author notes:** Correspondent author, Carlos Marcelino Rodriguez Lopez.

## Abstract

Enset (*Ensete ventricosum* (Welw.) Cheesman) is a drought tolerant, vegetatively propagated crop that was domesticated in Ethiopia. It is a staple food for more than 20 million people in Ethiopia. Despite its current importance and immense potential, enset is among the most genetically understudied and underexploited food crops. We collected 230 enset wild and cultivated accessions across the main enset producing regions in Ethiopia and applied amplified fragment length polymorphism and genotype by sequencing (GBS) methods to these accessions. Wild and cultivated accessions were clearly separated from each other, with 89 genes found to harbour SNPs that separated wild from cultivated accessions. Among these, 17 genes are thought to be involved in flower initiation and seed development. Among cultivated accessions, differentiation was mostly associated with geographical location and with proximity to wild populations. Our results indicate that vegetative propagation of elite clones has favoured capacity for vegetative growth at the expense of capacity for sexual reproduction. This is consistent with previous reports that cultivated enset tends to produce non-viable seeds and flowers less frequent than wild enset.

## Introduction

Plant domestication and breeding can alter and shrink genetic diversity (Miller & Schaal, 2006; Martínez-Ainsworth & Tenaillon, 2016). In some crop species, this entails a shift from sexual to vegetative propagation (Silvertown, 2008). Enset (*Ensete ventricosum* Welw.) Cheesman), often referred as false banana) is a hapaxanth diploid (2n=18) plant (Cheesman, 1947) that belongs to the Musaceae family (Shumbulo et al., 2012). In the wild, enset propagates by seed (Haile et al. 2014). The native distribution of wild enset encompasses the eastern coast Africa, from South Africa to Ethiopia, and extends west into the Congo (Borrell et al., 2019). In Ethiopia, which is considered to be the centre of origin of *E. ventricosum*, wild enset grows mainly along riversides and deep forest, extending into cultivated land and gardens in some regions (Olango *et al.*, 2015; Eshetae *et al.*, 2019).

Despite the wide distribution of wild enset, enset has been domesticated only in the Ethiopian highlands (Borrell et al., 2019; Heslop-Harrison et al., 2019) and it is now grown as a crop mainly in the southern and south-western parts of Ethiopia (Olango *et al.*, 2014; Guzzon & Müller, 2016). In these regions, cultivated enset is propagated vegetatively from suckers. Ethiopia maintains more than 600 accessions of cultivated enset *via* vegetative propagation (Harrison et al., 2014).

Due to its importance for food security in Ethiopia (Yemataw *et al.*, 2014; Guzzon & Müller, 2016; Yemata, 2020), enset has been called “the tree against hunger” (Brandt et al., 1997a). Enset is known for its high yield, drought tolerance, high shade potential, broad agro-ecological distribution and long storage capacity (Brandt et al., 1997b; Quinlan et al., 2014). Despite these positive features, enset has received little research attention (Borrell et al., 2019) and its genetic diversity is under threat from diseases such as bacterial wilt and from pressures associated with human population growth (Birmeta, 2004; Guzzon & Müller, 2016).

Genetic analysis of intraspecific variation in enset has mainly relied upon data for ‘anonymous’ molecular markers, such as amplified fragment length polymorphisms (AFLP) (Tsegaye & Struik, 2002), random amplified polymorphic DNA (RAPD) (Birmeta et al., 2004), inter simple sequence repeats (ISSR) (Tobiaw & Bekele, 2013) and microsatellites (simple sequence repeat (SSR) polymorphisms (Getachew *et al.*, 2014; Olango *et al.*, 2015; Gerura *et al.*, 2019). Given that enset is vegetatively propagated, genetic divergence among cultivars may be minimal (McKey et al., 2010) and could be difficult to detect using these marker types.

Here, we report on the application of both AFLP and next-generation sequencing (NGS) methods to 230 enset accessions (192 cultivated and 38 wild). Data collected using these methods were used to investigate population structure of cultivated and wild enset accessions and to identify signatures of selection and domestication within the enset genome. To our knowledge, this is the first application of NGS to a large number of accessions of wild and cultivated enset collected from a large geographic area.

## Results

### AFLP analysis

Based on the analysis of presence/absence data for 111 AFLP amplicons with lengths ranging from 51 bp to 350 bp, the heterozygosity and Shannon’s Index were higher for cultivated accessions (0.193±0.02 and 0.298±0.029) than for wild accessions (0.186±0.02 and 0.285±0.029). However, the average genetic distance between cultivated accessions was lower (0.026±0.002) than between wild accessions (0.047±0.007). The average percentage of polymorphic peaks for cultivated and wild accessions were 45.75% ± 3.25% and 41.7%±6.18 respectively.

Analysis of molecular variance (AMOVA) showed that that the majority (87-89%) of enset genetic variability is explained by within-region differences, while 11-13% can be attributed to variation between regions (Table 2). Principal coordinate analysis (PCoA) using AFLP markers showed that wild and cultivated enset accessions formed clusters with considerable overlapping of individuals from the two groups (Fig. 2a). Mantel test analysis showed significant correlation (r = 0.7; P< 0.0001) between genetic and geographic distances among cultivated and wild enset accessions (Supplementary Fig.1).

**Table 1:**
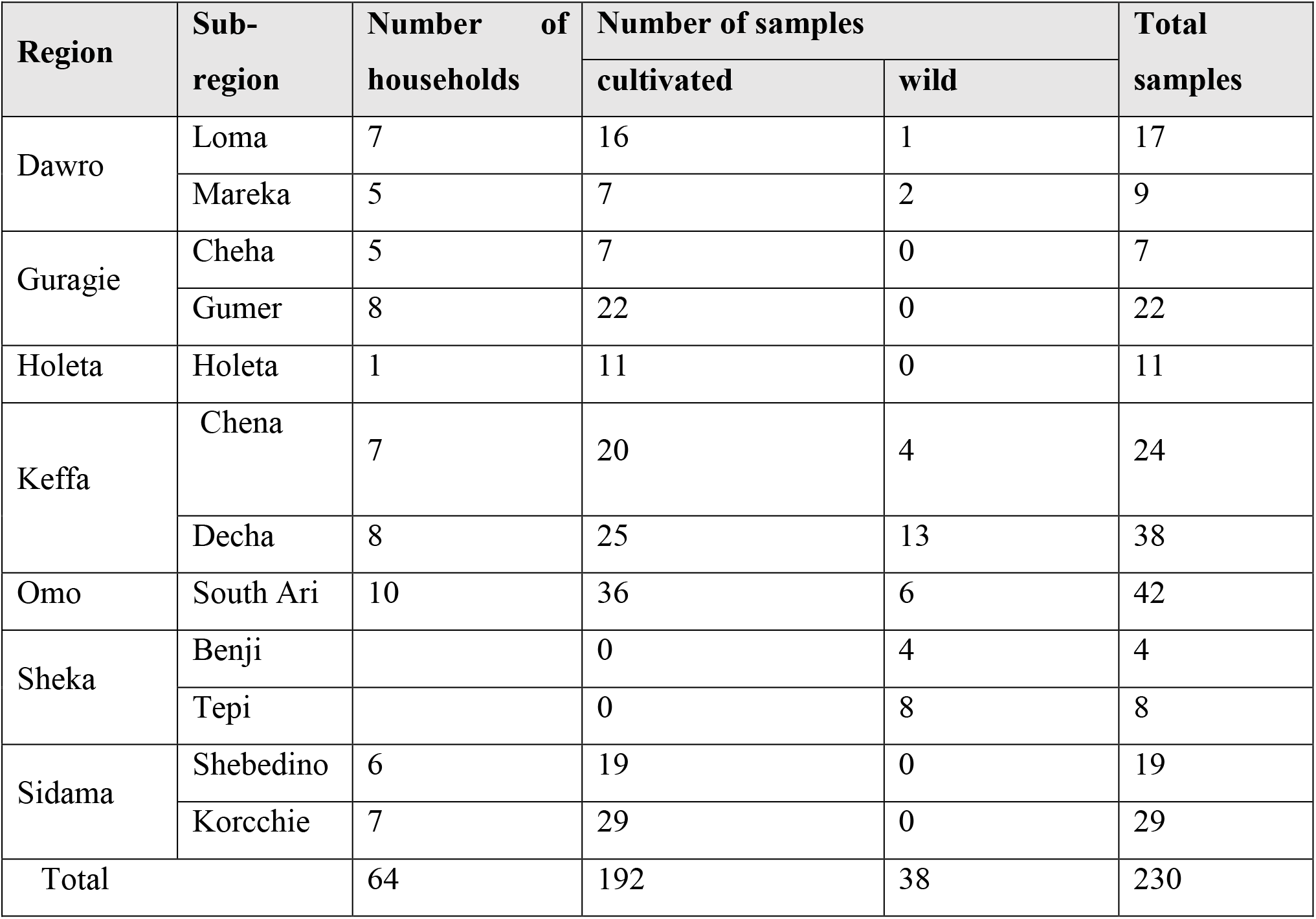
Summery of cultivated and wild enset accessions sampled from six regions in South and South Western Ethiopia.

| Region | Sub-region | Number of households | Number of samples |  | Total samples |
| --- | --- | --- | --- | --- | --- |
|  |  |  | cultivated | wild |  |
| Dawro | Loma | 7 | 16 | 1 | 17 |
|  | Mareka | 5 | 7 | 2 | 9 |
| Guragie | Cheha | 5 | 7 | 0 | 7 |
|  | Gumer | 8 | 22 | 0 | 22 |
| Holeta | Holeta | 1 | 11 | 0 | 11 |
| Keffa | Chena | 7 | 20 | 4 | 24 |
|  | Decha | 8 | 25 | 13 | 38 |
| Omo | South Ari | 10 | 36 | 6 | 42 |
| Sheka | Benji |  | 0 | 4 | 4 |
|  | Tepi |  | 0 | 8 | 8 |
| Sidama | Shebedino | 6 | 19 | 0 | 19 |
|  | Korcchie | 7 | 29 | 0 | 29 |
| Total |  | 64 | 192 | 38 | 230 |

**Table 2:** Analysis of Molecular Variance (AMOVA) using AFLP markers for 192 accessions of cultivated enset collected from seven regions

| Source | df | SS | MS | Est.Var. | % of variation | P-Value |
| --- | --- | --- | --- | --- | --- | --- |
| Among Populations | 5 | 173.320 | 34.664 | 0.915 | 13% | 0.0001 |
| Within Populations | 186 | 1175.024 | 6.317 | 6.317 | 87% |  |
| Total | 191 | 1348.344 |  | 7.232 | 100% |  |

df, degree of freedom; Est.Var, estimated variance; SS, sum of squares; %, the % of variance explained by within and across regions

**Fig.1.**
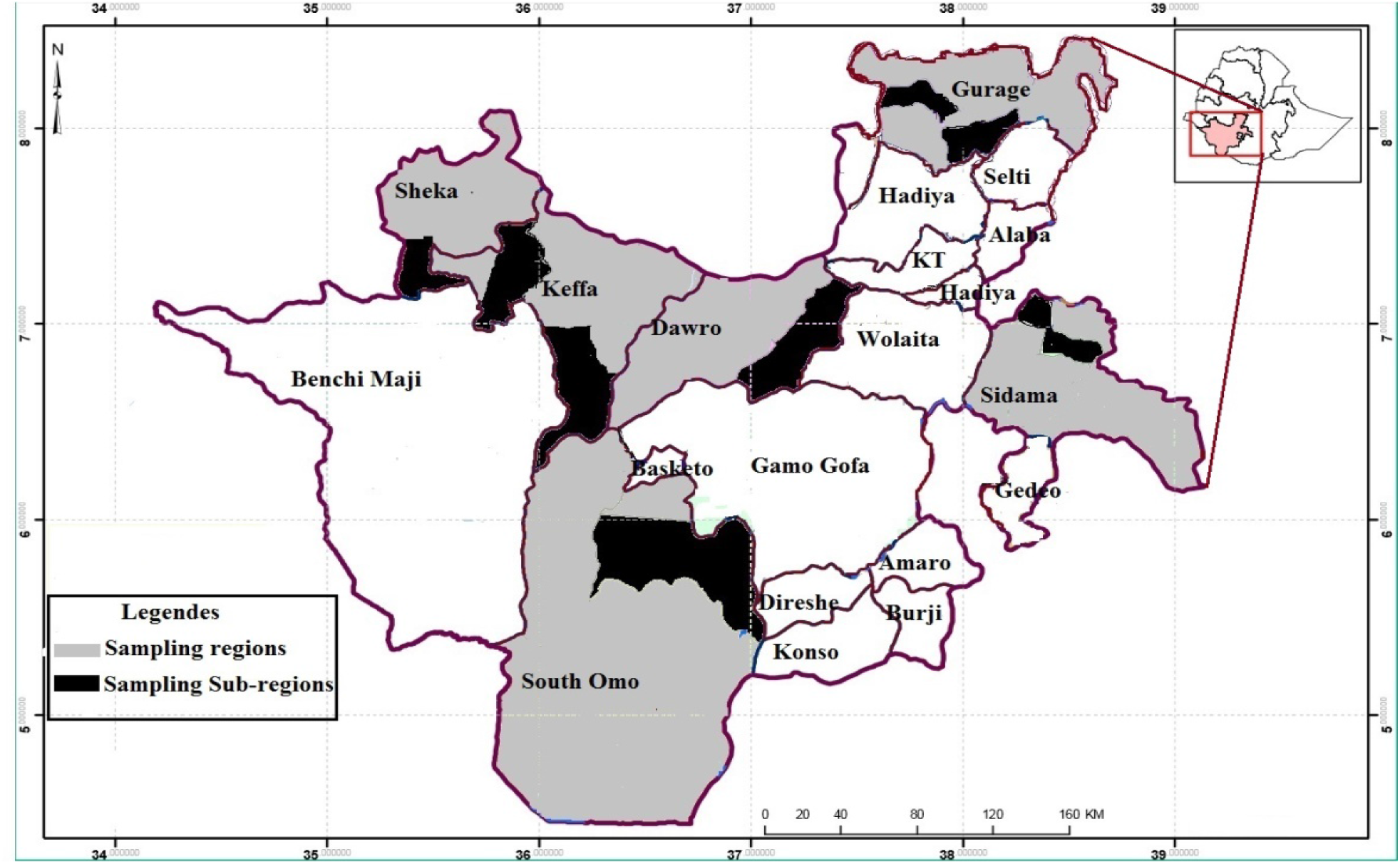
Sampling regions and sub-regions located in Southern Nation, Nationalities and People Region (SNNPR), Ethiopia.

**Fig.2.**
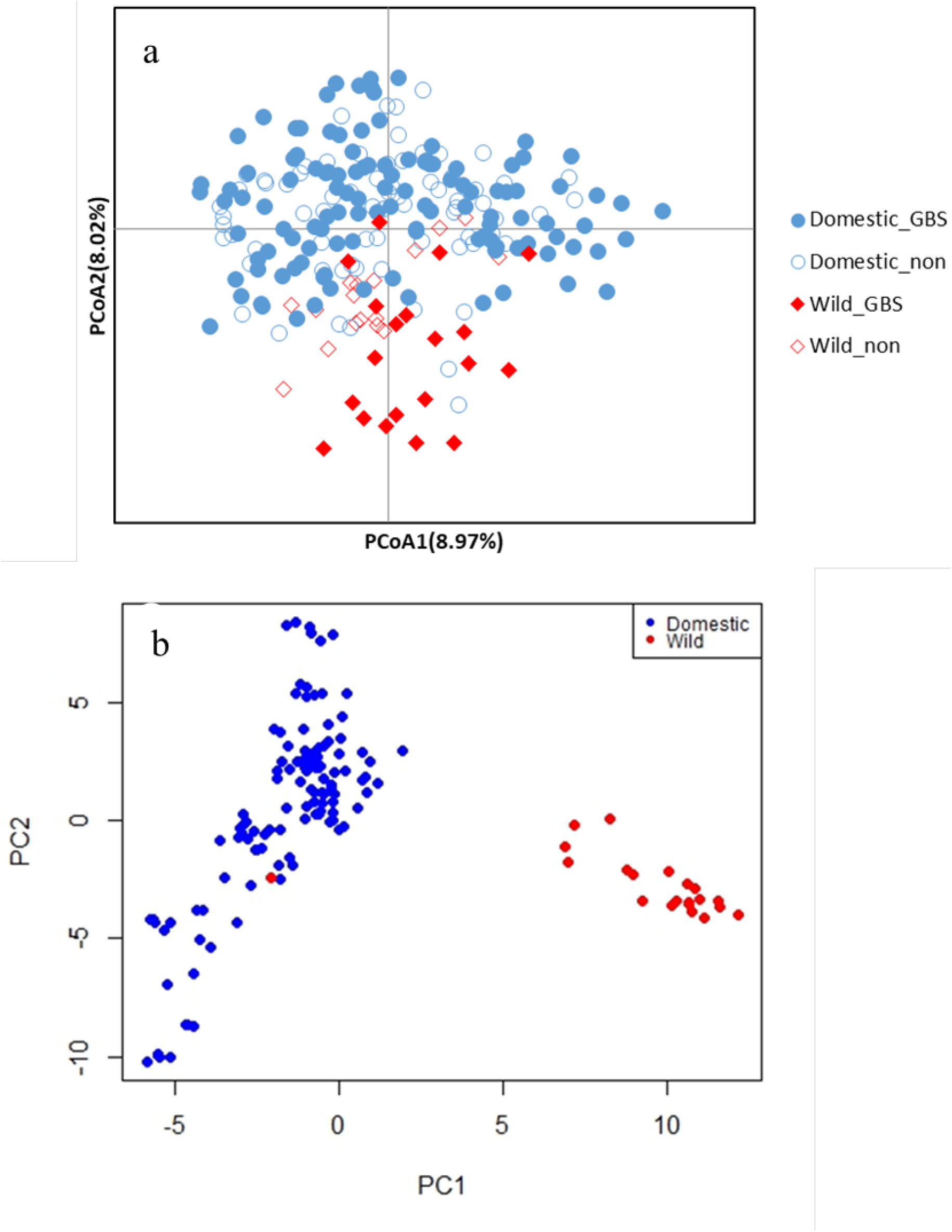
**a**) Principal Coordinate Analysis (PcoA) of 192 cultivated and 38 wild enset samples which are selected (141) and not selected (89) for GBS analysis. **b**) Principal Component Analysis (PCA) of 120 cultivated and 21 wild of enset accessions collected six top enset producing regions of Ethiopia, using GBS-based genome-wide SNPs 5169 SNP markers

### SNP discovery and analysis

Genotyping by sequencing of 149 (125 cultivated and 24 wild) enset accessions generated a total of 569,324,179 reads with 74 bp length and 50% of GC content. Eight samples were removed because of high SNP missing ratio, leaving 141 samples (120 cultivated and 21 wild) for analysis.

A total of 3,743,487 tags passed mapping criteria when physically mapped to the *Musa malaccensis* (wild banana) genome. This reference genome based SNP calling generated 22,884 SNPs showing locus coverage lower than 0.1 and minor allele frequency lower than 0.01. After filtering to remove SNPs with missing value greater than 20% and missing ratio greater than 30%, 5169 high quality SNPs remained. Of these, 4282 SNPs (83%) physically mapped to one of the 11 chromosomes of the *Musa malaccensis* genome and the remaining 887 SNPs were physically mapped to *Musa malaccensis* genome scaffolds (Table 3). The number SNPs per chromosome ranged from 251 in chromosome 2 to 465 in chromosome 4 (Table 3), with an average 389 SNPs per chromosome. The highest density of SNPs was detected on chromosome 4 (65.05 kb/SNP) and the lowest on chromosome 10 (91.5 kb/SNP). A/G transitions presented the highest frequency (29.06%) followed by C/T transitions (28.03%) and A/C transversions (11.30%) (Table 5).

**Table 3:** Total number of filtered and unfiltered SNP markers distributed across 11 chromosomes and chr_unknown (SNPs within contigs that had not been assigned to a chromosome in the assembly).

| Chromosome | Number SNP markers |  |
| --- | --- | --- |
|  | Unfiltered | Filtered |
| 1 | 1644 | 422 |
| 2 | 1262 | 251 |
| 3 | 1801 | 387 |
| 4 | 1978 | 462 |
| 5 | 1702 | 399 |
| 6 | 2056 | 417 |
| 7 | 1732 | 410 |
| 8 | 1955 | 442 |
| 9 | 1709 | 396 |
| 10 | 1708 | 368 |
| 11 | 1536 | 327 |
| ChrUn_random | 3800 | 887 |
| Total | 22883 | 5169 |

### Genetic relatedness and population structure of cultivated and wild enset accessions

PCA using the 5169 SNP markers indicated that all but one of the wild enset accessions clustered separately from the cultivated enset accessions (Fig.2b). UPGMA based phylogenetic tree (Fig.3) showed that the cultivated and wild enset accessions formed two clearly separated clades. The cultivated enset accessions formed multiple subclades within the cultivated enset population.

**Fig.3.**
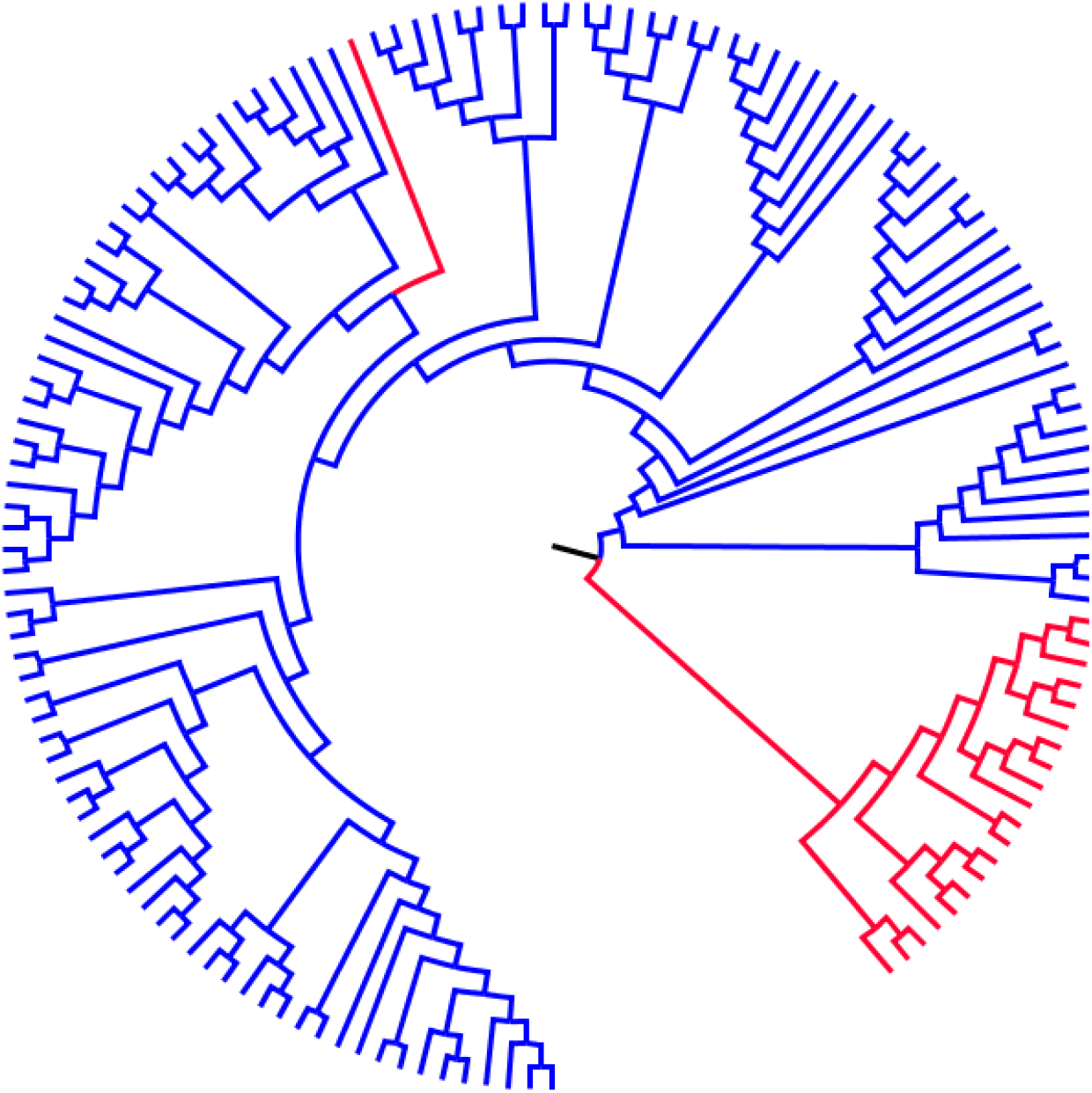
UPGMA phylogenetic tree of 120 cultivated (blue clades) and 21 wild (red clades) of enset accessions using GBS-based genome-wide SNPs 5169 SNP markers.

DAPC analysis showed a clear separation of enset accessions into three clusters (Fig.4a). Cluster 1 was comprised of 24 cultivated accessions and one wild accession. Among the 24 cultivated accessions in this clade, 17 accessions (71%) were collected from areas in which only cultivated enset was found. Cluster 2 contained 96 cultivated accessions, 67% of which were collected from areas which have both cultivated and wild enset accessions. Cluster 3 contained only wild enset accessions. STRUCTURE analysis with the Δ*K* method indicated the most informative number of subpopulations is two (*K* =2) (Fig. 5b). Individuals were considered part of a cluster when the probability of membership was 0.5 or greater. With *K*=2, 46 cultivated enset accessions clustered together with 20 wild enset accessions. With *K*=3, the pattern was similar (Fig.5a). At higher values of *K*, wild accessions continued to group together and the cultivated accessions grouped into three main clusters.

**Fig.4.**
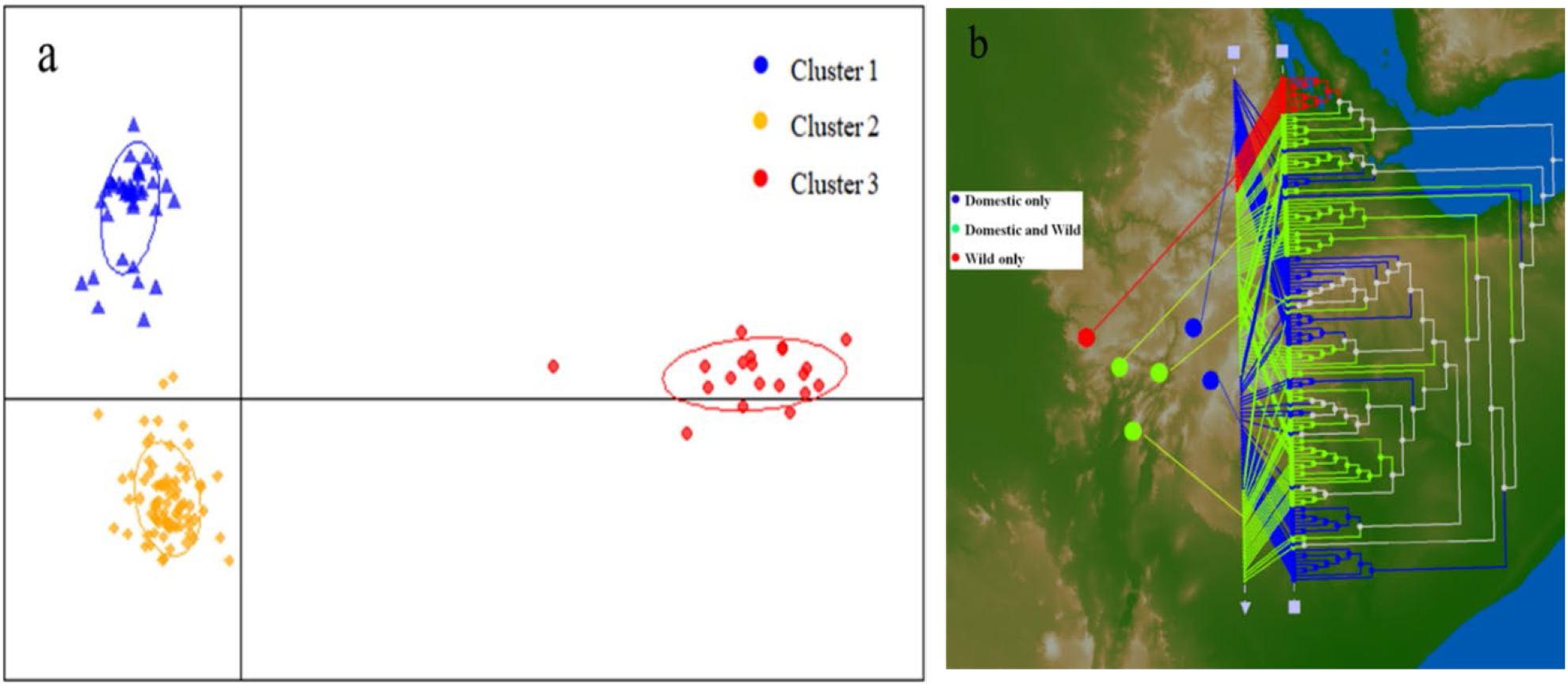
Genetic structure of 141 enset accessions using 5169 genome-wide SNP markers **a**) Population genetic structure using Discriminant Analysis of Principal Components (DAPC). b) GenGIS plot for the three clusters plotted with phylogenetic tree combined with the corresponding regions of collection. Samples were collected from different regions, regions with both domestic and wild, only domestic and only wild enset accessions.

**Fig.5.**
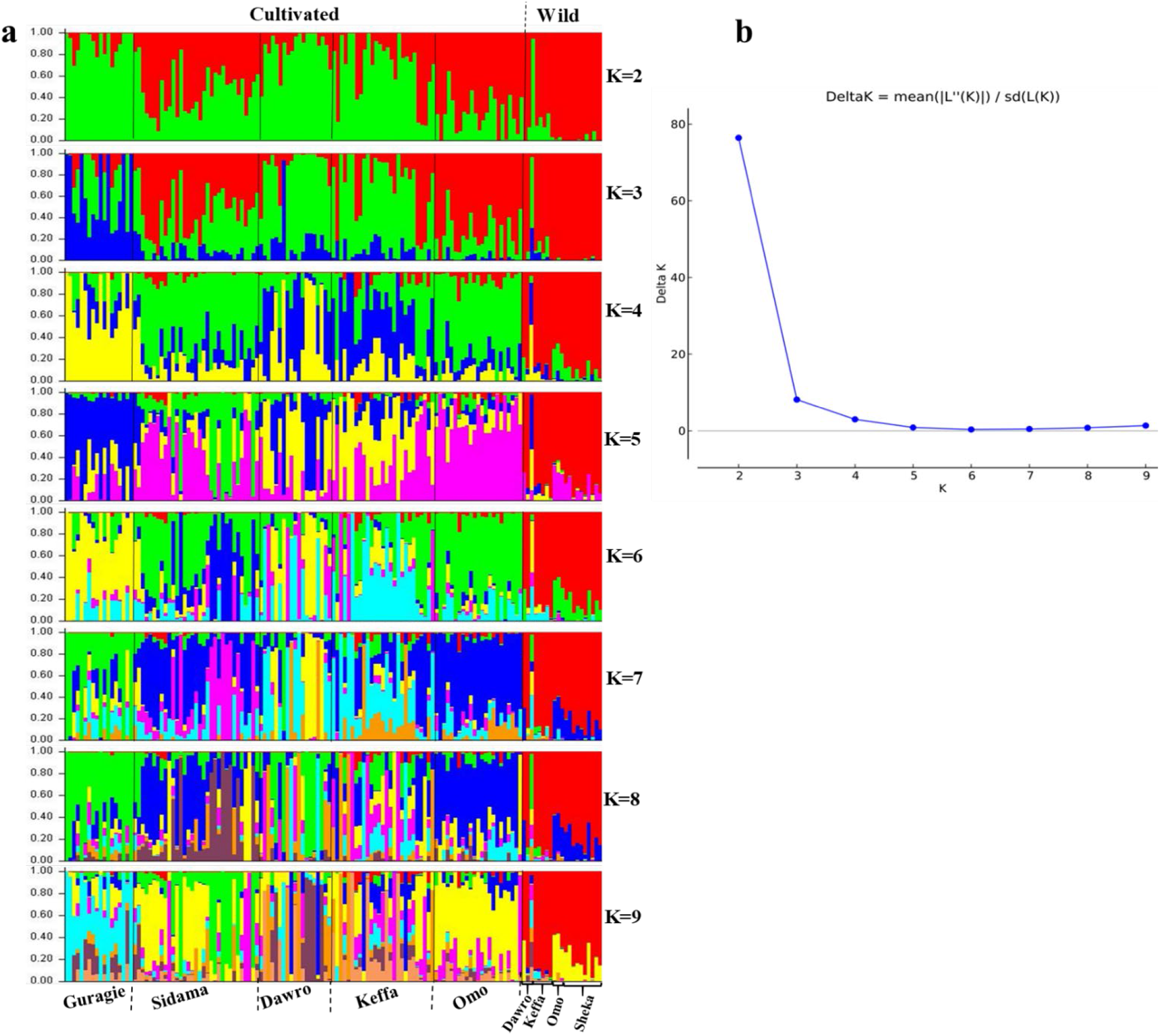
**a**) Estimated population structure of 141 cultivated and wild enset accessions analysed using the software STRUCTURE. Each accession is represented with vertical line, which is partitioned into coloured segments which represent the estimated membership fraction in the *K* clusters b) Evanno plot of *Delta K* calculated from K ranging from 2 to 9 (each K repeated 10 times) analysed using Structure-Harvester (Evanno *et al.*, 2005).

### Genetic diversity of cultivated and wild enset

The average PIC and gene diversity were similar for cultivated and wild accessions. Cultivated enset accessions exhibited higher heterozygosity than wild accessions, while the average major allele frequency was higher for cultivated than for wild accessions (Table 4). The average genetic distances and average F_*st*_ values among cultivated and wild accessions were 0.33±0.001(SE) and 0.11±0.005 (SE), respectively.

**Table 4:** Genetic diversity analysis of cultivated and wild enset accessions collected from six major enset producing regions of Ethiopia, analysed using 5169 GBS-based SNP markers

| Category | Sample Size | Major allele frequency | Gene Diversity | Heterozygosity | PIC |
| --- | --- | --- | --- | --- | --- |
| Cultivated | 120 | 0.71±0.002 | 0.40±0.003 | 0.26±0.003 | 0.35±0.003 |
| Wild | 21 | 0.68±0.002 | 0.42±0.002 | 0.21±0.002 | 0.37±0.002 |

**Table 5:** Number and proportion of nucleotide combinations of the 5169 SNP markers

| Allele | Number | Proportion (%) |
| --- | --- | --- |
| A/C | 567 | 10.97 |
| A/G | 1502 | 29.06 |
| A/T | 431 | 8.34 |
| C/G | 504 | 9.75 |
| C/T | 1449 | 28.03 |
| G/T | 580 | 11.22 |
| multiple allele | 136 | 2.63 |
| Total | 5169 | 100 |

Analysis of genome-regional patterns of nucleotide diversity using 500 kb non-overlapping sliding windows showed that the average nucleotide diversity was higher in wild enset accessions (0.32±0.005 (SE)) than in cultivated enset accessions (0.27±0.006 (SE)) (Fig.6). Calculation of the degree of diversification (F_*st*_) between cultivated and wild enset accessions identified a total of 29 genomic subregions with high degree of diversification (F_*st*_>0.2) and 76 genomic subregions with low F_*st*_ (F_*st*_<0.02) (Fig.6). Chromosomes 3, 5 and 10 presented the highest number of genomic subregions with high F_st_. On the other hand, chromosome 1 presented the highest number of low F_*st*_ genomic subregions (11 subregions), while chromosomes 2, 3 and 5 showed the lowest number of low F_st_ genomic subregions (4 subregions).

**Fig.6.**
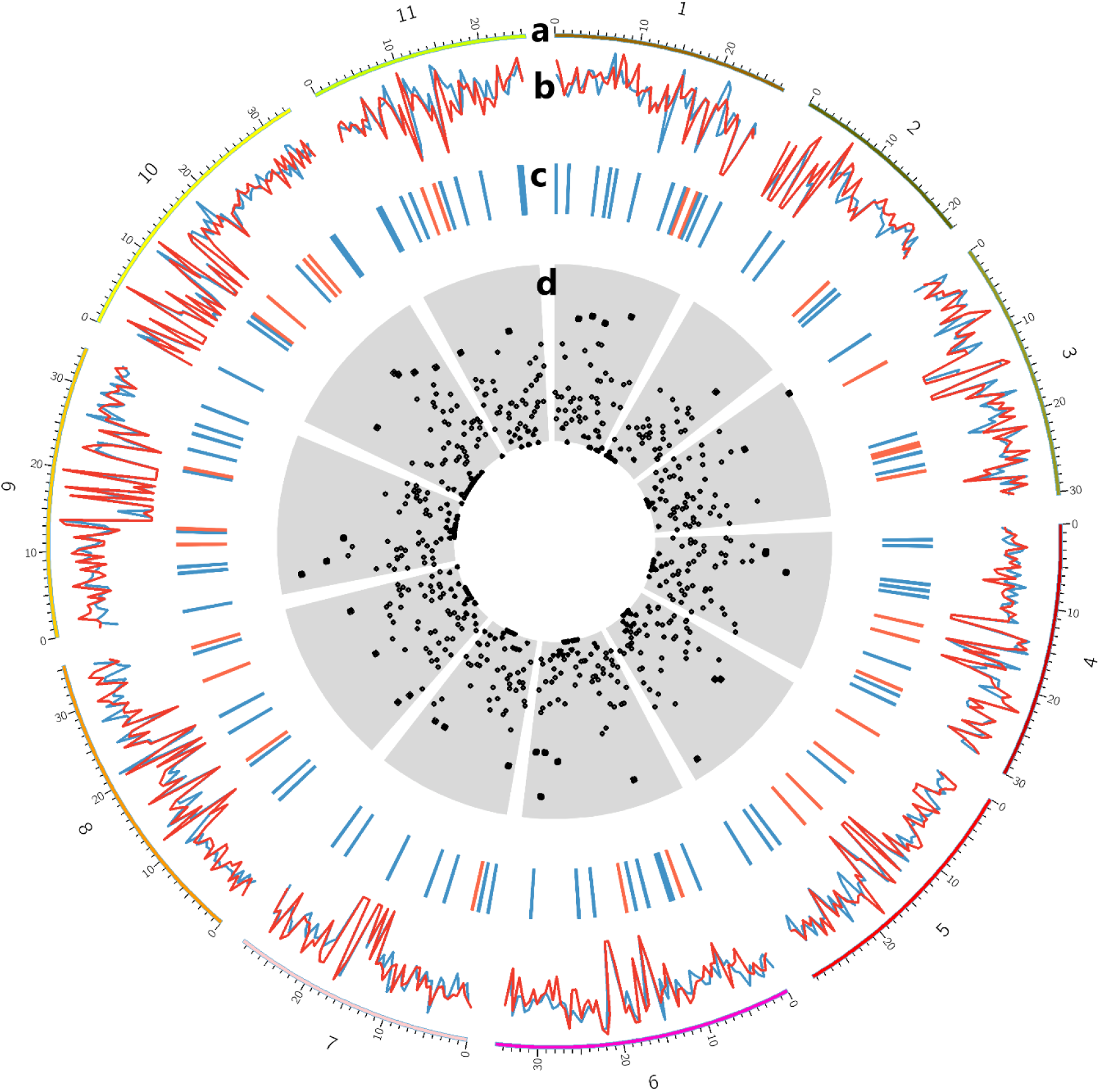
Summary of genetic diversity and genetic differentiation of cultivated and wild enset accessions measured within 500kb sliding window drawn using circos plot. **a**) The 11 Chromosomes (Mb) portrayed along the perimeter of each circle, **b**) Genetic diversity of cultivated (blue) and wild (red) enset accessions, genetic diversity for each sliding window was calculated nucleotide diversity divided by number of markers. **c**) F_*st*_ less than 0.02 (red) and greater than 0.2 (blue), **d**) total count of SNP markers per window, dots near the centre represent a low number of SNPs and the dots further out represent high numbers of SNPs.

### Genomic regions under selection pressure

The genome scan approach (LOSITAN-based *F_st_*-outlier detection method) implemented in this study identified 158 (2.56%) SNPs which frequency was significantly different between cultivated and wild enset populations which are dispersed throughout the 11 chromosomes of the wild banana reference genome (Fig. 7). Chromosome 3 and 10 harbour highest number of outlier SNPs (16 outlier SNPs each). Chromosome 1 contains the lowest number of outlier SNPs (4 SNPs), despite containing the highest number of SNP markers. Mapping of outlier SNPs to the reference genome (*Musa acuminata subsp. malaccensis*) genome identified 89 genes containing one or more SNPs within their protein coding region (Supplementary Table 4). Of these, 19 genes were found to be associated to sexual reproduction traits, i.e. flowering (8 genes), seed development and germination (9 genes) or domestication (2 genes) (Supplementary Table 6). The function of these genes were annotated based on comparative genomics (one gene), deduced from protein containing domains as putative function (five genes) and experimentally validated (14 genes) in other plants such as Arabidopsis thaliana, rice, soybean and tomato.

**Fig.7.**
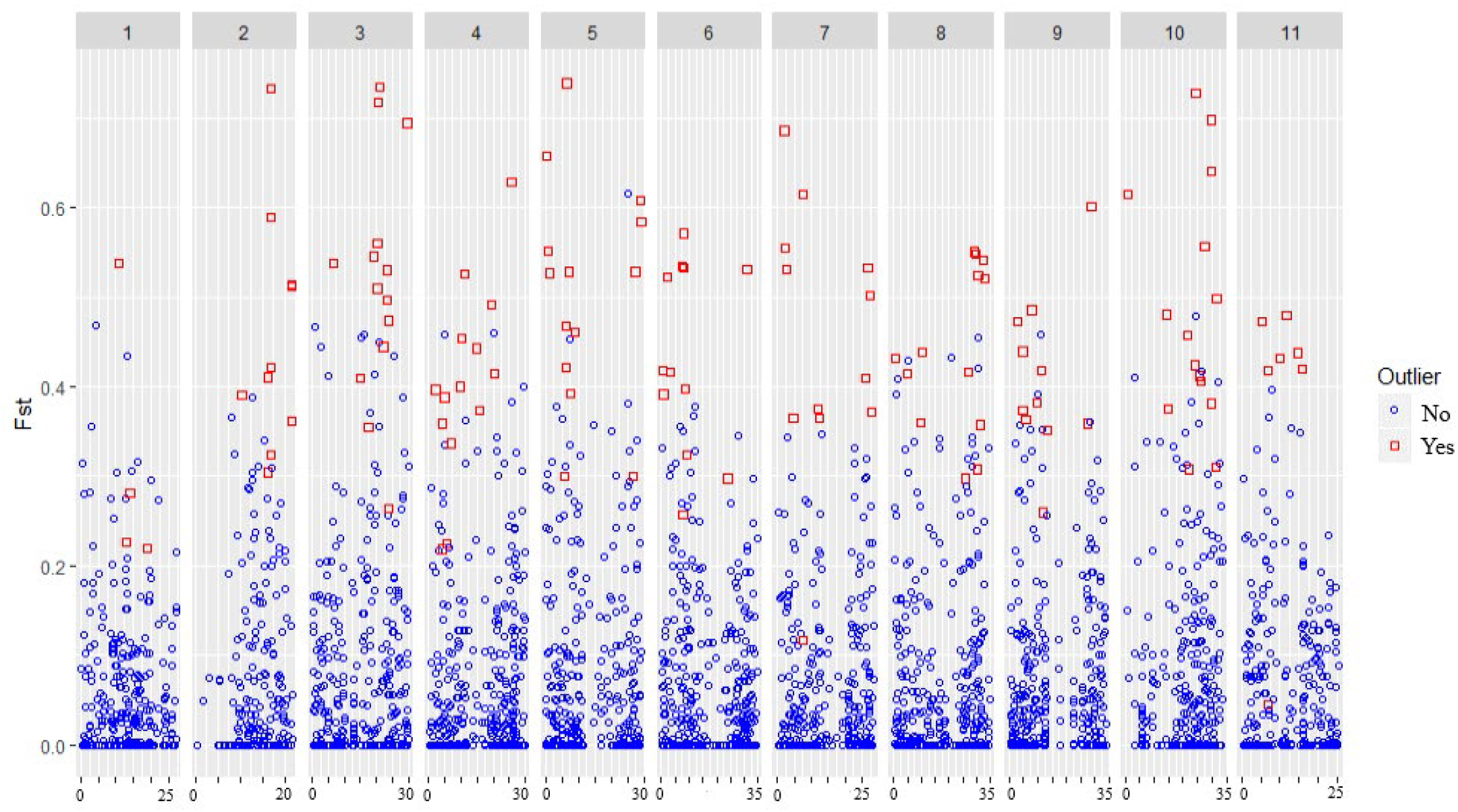
F_*st*_ values of 5169 SNP loci, displayed according to their genomic positions within 5 Mb intervals on the 11 chromosomes.

## Discussion

### Genetic diversity of enset in Ethiopia

The results presented here indicate that cultivated and wild enset accessions exhibit similar gene diversity and polymorphic information content (PIC) (Table 4). This is similar to what has been reported based on SSR marker analysis of enset genetic diversity (Gerura *et al.*, 2019), but differs from what has been reported by Olango *et al.* (2015), which reported that higher gene diversity in cultivated enset population (0.59) than wild enset population (0.40), but similar hetrozygosity for cultivated and wild enset populatons (0.5). The genetic diversity for both cultivated and wild enset reported in the current study (Table 4) is lower than previous enset genetic diversity studies conducted using SSR makers (Getachew *et al.*, 2014; *Olango et al., 2015*; Gerura *et al.*, 2019). These differences and discrepancies might be due to the nature of the different types of markers used. SSRs and microsatellite are multi-allelic and are more polymorphic than SNP markers, which are usually bi-allelic. The genetic diversity detected here for enset is higher than what has been reported for some other vegetatively propagated plants such as Cassava (de Albuquerque *et al.*, 2018; Kamanda *et al.*, 2020), and out-crossing plants such as sunflower (Mandel et al., 2011) but lower what has been reported for Japonica rice (Becerra et al., 2017).

Our observations that cultivated ensets exhibited higher heterozygosity and Shannon's Index than wild enset resemble what have been reported for enset based on SSR markers study (Gerura *et al.*, 2019) and for other plant species, including *Camellia sinensis* (Yang et al., 2016) and *C. taliensis* (Zhao et al., 2014). The high heterozygosity of cultivated enset might be due to vegetative propagation maintaining heterozygosity across clonal generations. In addition, the wild enset habitat has been sharply declining in Ethiopia because of population growth and deforestation (Birmeta et al., 2004; Olango et al., 2015). This reduction in effective population size might have contributed to the observed lower heterozygosity due to the increase of chances of inbreeding in wild enset populations. However, genetic distances were greater among wild accessions than among cultivated accessions, possibly because wild populations remained isolated by distance or geographical barriers (Tobiaw & Bekele, 2013), while cultivated materials were more readily transferred between regions through regular long-distance accessions exchange between farmers (Brimata et al.,2020). Limited genetic distances among cultivated enset accessions could also be due to recent separation (fragmentation) of the varieties, without sufficient evolutionary time to generate variation (Burgos-Hernández et al., 2013).

### Population structure and genetic relationship between cultivated and wild enset accessions in Ethiopia

PCA and phylogenetic analysis revealed that cultivated and wild enset accessions separated into genetically distinct clusters despite being morphologically similar members of the same taxonomic species. This indicates that cultivated enset accessions has been domesticated from a limited number of wild progenitors (Brimeta et al.,2004). It is also possible that currently cultivated enset and the currently available wild enset in Ethiopia originated from different ancestral materials. Bayesian clustering program STRUCTURE indicated that cultivated population grouped into three clusters and this is similar to previous SSR marker based enset genetic diversity study (Olango *et al.*, 2015). The genetic sub-clustering within cultivated enset corresponds with geographical distance between the accessions and proximity to wild enset. Cultivated enset accessions collected from areas where wild enset grows showed higher admixture and weaker clustering than those collected from areas where wild enset does not grow. Some cultivated accessions clustered with wild accessions, possibly indicating recent introgression of wild enset into farming systems. In the Omo region, particularly in the Ari sub-region, wild enset growing in gardens have been adopted by farmers as a cultivated crop and propagated (Shigeta, 1990; Hildebrand, 2001a). Thus, multiple domestication events and/or frequent introgression from wild enset could explain the high genetic diversity and overlapping spatial distributions of wild and cultivated enset (Borrell et al., 2019). Mantel test showed significant correlation (r=0.7, P=0.0001) between genetic and geographic distances separating wild and cultivated populations. Our results support Olango *et al* (2015) and Brimeta *et al* (2004) reports of a limited possibility of gene flow due to the natural distribution of wild enset and farming management.

### Loci under selection signature

Improved understanding of the genetic adaption of enset could facilitate genetic improvement. F_*st*_ outlier tests for detecting extreme allele frequency differentiation can detect genomic regions that have evolved under adaptation and selection (Beaumont & Balding, 2004; Lotterhos & Whitlock, 2014). Here, application of an F_*st*_ outlier test identified 158 outlier SNP markers that show significant (P<0.01) genetic differentiation between cultivated and wild enset. Mapping of these outlier markers to the diploid banana genome led to the identification of 89 genes under selection during enset domestication. 19.1% of these genes were found to be related to the regulation of flowering (Zhong & Ye, 2004; Gunesekera *et al.*, 2007; *Ishida et al., 2008*; Uehara *et al.*, 2019; Kang *et al.*, 2020), or seed development (Swain *et al.*, 2005; Wen *et al.*, 2008; Li & Li, 2014; Ma *et al.*, 2018). Interestingly, another 2.3% of the genes found to be under selection have been previously associated to domestication in other species (Chakrabarti *et al.*, 2013; Jan *et al.*, 2013; Li *et al.*, 2017). Flowering and seed development are important characteristics that differentiate cultivated and wild enset (Borrell et al., 2019). Wild enset flowers more frequently and has larger flowers (mean basal girth 186cm) than cultivated enset (mean basal girth 106cm) (Shigeta, 1990; Hildebrand, 2001b). Wild enset is highly prolific, producing thousands of large (about 12 mm diameter) hard black seeds, while cultivated enset plants bear fewer seeds, which are small (3 mm), soft, pale and incompletely developed (Hildebrand, 2001b). It has been previously suggested that these traits could be due to reduced fitness resulting from a subsequent selection and domestication bottleneck (Heslop-Harrison et al., 2019). The proportion of genes found to be under selection in cultivated enset, certainly indicate that selection associated to domestication could be the driver of those traits.

In addition, the calculation of degree of diversification (F_*st*_) between cultivated and wild enset accessions enabled the identification of genomic subregions (500 kb non-overlapping) with high (F_*st*_ > 0.2) degree of diversification (Fig 6). Genomic subregions with high F_*st*_ may contain or associated to potential genes that are related to plant domestication and adaptions, and provide an indication of the functional genes involved (Lam *et al.*, 2010). In the current study, F_*st*_ outlier based scan for candidate genes under putative selection and adaptation has found promising results and is an important step forward to further studies on gene mapping and identification, and designing enset breeding program. to better understand enset genome and use our result, it is necessary to conduct further gene mapping and genome-wide association studies with large sample size covering all the enset producing regions of Ethiopia.

## Materials and methods

### Study area

Samples were collected from six of the major enset producing regions in Ethiopia: Dawro, Guragie, Keffa, South Omo, Sheka and Sidama (Fig.1). Within each of these regions, samples were collected from subregions and from two or three districts within each subregion (Table 1). Within each district, samples of domesticated enset were collected from five to ten households, selected based on recommendations from local agricultural extension experts. Samples of wild enset were collected around farming areas, along riversides and in deep forests. For each sampling location, latitude and longitude (degrees, minutes, seconds) were collected using GPS essentials mobile app (https://downloads.tomsguide.com/GPS-Essentials,0301-49666.html) and then transformed to standard Universal Tranverse Mercator coordinates (UTM) using a geographic unit converter (http://www.rcn.montana.edu/Resources/Converter.aspx) (Supplementary Table 1).

### Sample collection and DNA extraction

Leaf samples were collected from 230 (192 cultivated and 38 wild) enset plants, each of which was between one and two years old (based on the farmer’s information) (Supplementary Table 1). Each sample consisted of a 5 cm * 5 cm fragments of the leaf blade of the most recently unfurled leaf. Each sample was placed in a 50 ml tube and stored on ice during transportation, then stored at −80 °C until DNA extraction. Each subsample (80-90 mg) were milled using a mortar and pestle immersed in liquid nitrogen. DNA extractions were performed using DNeasy Plant Mini Kits (Qiagene Inc.) according to the manufacturer’s instructions. DNA concentration was measured using the QuantiFluor^(R)^dsDNA System^(a)^ (Promega, USA) following manufacturer’s instructions, then adjusted to 20 ng/μl using molecular biology grade water (Sigma).

### Amplified Fragment Length Polymorphism (AFLP) preparation and analysis

AFLP reactions (Vos *et al.*, 1995) were performed for all 230 samples using a modification of the protocol described by López *et al.* (2012). Briefly, samples containing 55 ng of genomic DNA were enzymatically digested in a 12.5 reaction volume containing *Mse*I, *EcoR*I (NEB) and ligated to *Mse*I and *EcoR*I adaptors (Supplementary Table 4) at 37° C for 2 h in a T100™ Thermal cycler (*Bio-Rad* Laboratories, Hercules, CA). Success of the digestion/ligation reaction was confirmed on 1.5% of agarose gel electrophoresis. Pre-selective PCR amplification was carried out using primers containing a 3’ selective nucleotide (i.e., *EcoR*I=A and *Mse*I=C). Selective amplification was then conducted using a primer combination with three selective nucleotides at the 3’ ends (*EcoR*I =*ACG*) and *Mse*I *=CAA*). Selective bases were chosen according to previous work on enset (Negash et al., 2002). PCR products were separated using Applied Biosystems 3130/3130xl Genetic Analysers (Applied Biosystems Life Technologies).

AFLP profiles were analysed using GeneMapper® Software v4.0. Clear and unambiguous polymorphisms were considered and were scored on a presence/absence basis for each marker. Clearly polymorphic peaks were verified manually and scored as present (1) or absent (0) for each sample. The level of AFLP polymorphism and genetic diversity across enset accessions were examined using GenAlEx 6.502 (Peakall & Smouse, 2012) based on average band frequency, Nei’s unbiased genetic distance, principal coordinate analysis (PCoA) and analysis of molecular variance (AMOVA). To examine possibility of gene flow between cultivated and wild enset accessions, the correlation between genetic distance (Φ_PT_) and geographic distance (km) was estimated using a Mantel test (Mantel, 1967) implemented in GenAlex using 10,000 random permutations.

### Genotyping by sequencing (GBS) preparation and analysis

Genotyping-by-sequencing was conducted for 149 enset samples (125 domestic and 24 wild; Supplementary Table 2) that were selected to capture the genetic diversity shown by AFLP. The GBS library preparation was carried out as described by Xie *et al.* (2017) including a water negative control as described by Konate *et al.* (2018). The DNA concentration of each individual library was normalized to 5 ng/μl. Two pooled libraries were created, each by pooling the individual libraries from 75 uniquely barcoded samples (25 ng per sample) (Supplementary Table 2). Each pooled library was then amplified in 10 PCR reactions, each containing 10 μl of digested/ligated DNA library, 12.5 μl of NEB MasterMix, 2 μl of 10 μM forward and reverse Illumina_PE primers (Supplementary Table 4) and 0.5 μl of molecular biology grade water (Sigma). The amplification reaction was carried using a T1000 Thermocycler at 95°C for 30 s, 16 cycles of (95°C for 30 s, 62°C for 20 s, 68°C for 30 s) and 72°C for 5 min. Amplification products were pooled together and cleaned using AMPure XP beads (Beckman Coulter, Australia) (1:1 ratio) to remove excess primers and unremoved adaptors. Libraries were sequenced using an Illumina NextSeq High Output 75 bp single-end run (Illumina 1.9 Inc., San Diego, CA, United States) at the Australian Genome Research Facility (AGRF, Adelaide, SA, Australia).

### GBS SNP calling

SNP calling was performed using two pipelines: *de novo*-based (reference genome independent) TASSEL-UNEAK pipeline (Lu et al., 2013) and the reference-based TASSEL-GBS pipeline (Glaubitz *et al.*, 2014; Torkamaneh *et al.*, 2016). Only sequences containing identical matches to the barcodes followed by the expected sequence of three nucleotides remaining from a *Msp*I cut-site (5’-CGG-3’) were selected for the identification of SNPs. FASTQ files containing barcoded sequence reads were demultiplexed using unique barcodes for each sample and trimmed to 64 bp (not including the barcodes). Identical sequence reads were collapsed into tags and sequencing tags from the four NextSeq Illumina sequencing lanes were merged to form one master tag. These sequence tags were mapped to the wild (diploid) banana (*Musa acuminata ssp. malaccensis*) genome sequence (D’hont et al., 2012) to deduce their genomic position. Tags with single base pair mismatches between samples were considered as SNPs and were generated in Hapmap format.

### Genetic diversity and population structure analysis

Genetic diversity and genetic differentiation (F_*st*_) were calculated using PopGenome R package (Pfeifer et al., 2014). Heterozygosity (the proportion of heterozygous individuals in the population), gene diversity (expected heterozygosity) and polymorphic information content (PIC) were calculated using Power Marker V3.25 (Liu & Muse, 2005). To examine the relationship between cultivated and wild enset accessions, PCA plots and phylogenetic tree (UPGMA) were built using TASSEL 5 (Bradbury et al., 2007). GenGIS (Parks et al., 2009) was used to display the phylogenetic tree with the geographic regions of sample collection.

Population structure was analysed using descriptive analysis of principal components (DAPC) (Jombart et al., 2010) and STRUCTURE (Pritchard et al., 2000). The software STRUCTURE was used to analyse the hierarchical population structure by setting the length of the burn-in period to 50,000 iterations and number of the MCMC replications after burn-in to 50,000. Between two to nine population clusters (K) were considered, with 10 iterations conducted for each *K*-value. The best *K*-value was determined using Structure Harvester (Earl, 2012) based on delta *K* (Δ*K*) (Evanno et al., 2005) and maximum log likelihood L(*K*) (Rosenberg *et al.*, 2001)

Genome-wide nucleotide diversity (average pairwise nucleotide differences) and population differentiation (F_*st*_) within and between wild and cultivated populations were calculated using a 500 kb non-overlapping sliding window. To obtain genetic diversity per window, nucleotide diversity was divided by number of SNPs per sliding window. These statistics were calculated using R package PopGenome (Pfeifer *et al.*, 2014) and plotted using Circos (Krzywinski et al., 2009) to visualize the pattern of genetic diversity across the whole enset genome.

To detect loci under selection during enset domestication and adaptation, the FDIST2 method adopted by Beaumont and Nichols (1996) was applied using lositan software (Antao *et al.*, 2008). F_*st*_ value was calculated for each SNP using allele frequencies conditional on expected heterozygosity (*He*), and P-values for each SNP were calculated. SNPs within tags assigned to one of the wild banana chromosomes were used to identify F_*st*_ outliers. F_*st*_ outlier analysis was carried out with 50,000 interactions at 99% confidence interval. Then we searched for genes containing these outlier SNPs in the wild banana genome to identify potential genes under selection during enset domestication using magrittr R package (Bache & Wickham, 2014) and generated gene ID. The putative function of these genes were searched using UNIPROT database (https://www.uniprot.org/) based on the gene ID.

## Acknowledgments

This study was conducted as part of a biosafety capacity-building project for sub-Saharan Africa implemented by the international centre for genetic engineering and biotechnology (ICGEB). This report is based on research funded by the Bill & Melinda Gates Foundation. The findings and conclusions contained within are those of the authors and do not necessarily reflect positions or policies of the Bill & Melinda Gates Foundation nor the ICGEB. Dr Lopez is currently partially supported by the National Institute of Food and Agriculture, U.S. Department of Agriculture, Hatch Program number 2352987000.

---

**Supplementary Fig.1.**
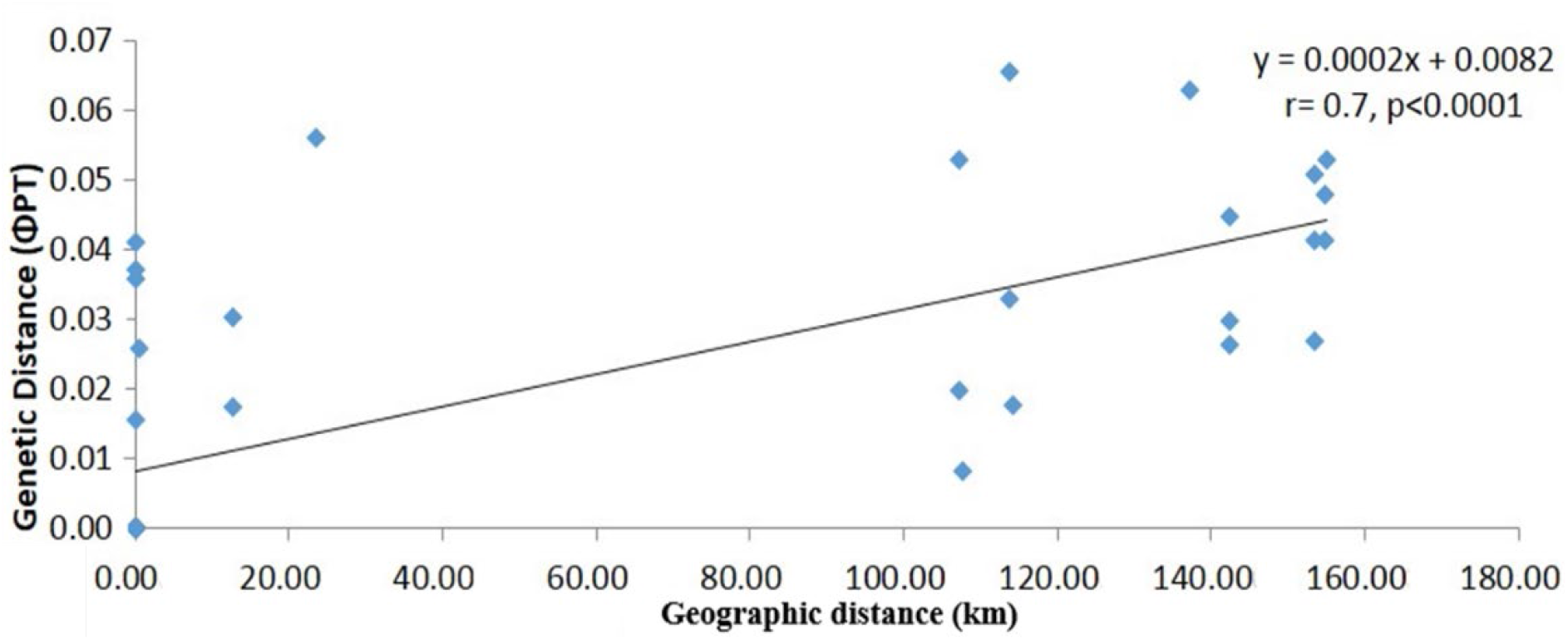
Mantel test to estimate correlation between genetic (ΦPT) measured using AFLP markers and geographic distance (Km) of cultivated and wild enset samples, including the regression formula, accuracy (r) and significance test (P).

## References

Antao T, Lopes A, Lopes RJ, Beja-Pereira A, Luikart G. 2008. LOSITAN: a workbench to detect molecular adaptation based on a F ST-outlier method. BMC bioinformatics 9(1): 323.

Bache SM, Wickham H. 2014. magrittr: a forward-pipe operator for R. R package version 1(1).

Beaumont MA, Balding DJ. 2004. Identifying adaptive genetic divergence among populations from genome scans. Molecular Ecology 13(4): 969–980.

Beaumont MA, Nichols RA. 1996. Evaluating loci for use in the genetic analysis of population structure. Proceedings of the Royal Society of London. Series B: Biological Sciences 263(1377): 1619–1626.

Becerra V, Paredes M, Ferreira ME, Gutiérrez E, Díaz LM. 2017. Assessment of the genetic diversity and population structure in temperate japonica rice germplasm used in breeding in Chile, with SSR markers. Chilean journal of agricultural research 77(1): 15–26.

Birmeta G. 2004. *Genetic variability and biotechnological studies for the conservation and improvement of Ensete ventricosum*. Alnarp: Sveriges lantbruksuniv., Acta Universitatis Agriculturae Sueciae.

Birmeta G, Nybom H, Bekele E. 2004. Distinction between wild and cultivated enset (Ensete ventricosum) gene pools in Ethiopia using RAPD markers. Hereditas 140(2): 139–148.

Borrell JS, Biswas MK, Goodwin M, Blomme G, Schwarzacher T, Heslop-Harrison J, Wendawek AM, Berhanu A, Kallow S, Janssens S. 2019. Enset in Ethiopia: a poorly characterized but resilient starch staple. Annals of botany 123(5): 747–766.

Bradbury PJ, Zhang Z, Kroon DE, Casstevens TM, Ramdoss Y, Buckler ES. 2007. TASSEL: software for association mapping of complex traits in diverse samples. Bioinformatics 23(19): 2633–2635.

Brandt SA, Spring A, Hiebsch C, McCabe JT, Tabogie E, Diro M, Wolde-Michael G, Yntiso G, Shigeta M, Tesfaye S 1997a. The “Tree Against Hunger”: Enset-Based Agricultural System in Ethiopia: American Association for the Advancement of Science Washington, DC, USA.

Brandt SA, Spring A, Hiebsch C, McCabe JT, Tabogie E, Diro M, Wolde-Michael G, Yntiso G, Shigeta M, Tesfaye S. 1997b. The tree against hunger. Enset-based agricultural systems in Ethiopia American Association for the Advancement of Science, Washington DC.

Burgos-Hernández M, Hernández DG, Castillo-Campos G. 2013. Genetic diversity and population genetic structure of wild banana Musa ornata (Musaceae) in Mexico. Plant systematics and evolution 299(10): 1899–1910.

Chakrabarti M, Zhang N, Sauvage C, Muños S, Blanca J, Cañizares J, Diez MJ, Schneider R, Mazourek M, McClead J. 2013. A cytochrome P450 regulates a domestication trait in cultivated tomato. Proceedings of the National Academy of Sciences 110(42): 17125–17130.

Cheesman E. 1947. Classification of the Bananas: The Genus Musa L. Kew Bulletin: 106–117.

D’hont A, Denoeud F, Aury J-M, Baurens F-C, Carreel F, Garsmeur O, Noel B, Bocs S, Droc G, Rouard M. 2012. The banana (Musa acuminata) genome and the evolution of monocotyledonous plants. Nature 488(7410): 213.

de Albuquerque HYG, Carmo CDd, Brito AC, Oliveira EJd. 2018. Genetic diversity of Manihot esculenta Crantz germplasm based on single‐nucleotide polymorphism markers. Annals of applied biology 173(3): 271–284.

Earl DA. 2012. STRUCTURE HARVESTER: a website and program for visualizing STRUCTURE output and implementing the Evanno method. Conservation genetics resources 4(2): 359–361.

Eshetae MA, Hailu BT, Demissew S. 2019. Spatial characterization and distribution modelling of Ensete ventricosum (wild and cultivated) in Ethiopia. Geocarto International: 1–16.

Evanno G, Regnaut S, Goudet J. 2005. Detecting the number of clusters of individuals using the software STRUCTURE: a simulation study. Molecular Ecology 14(8): 2611–2620.

Gerura FN, Meressa BH, Martina K, Tesfaye A, Olango TM, Nasser Y. 2019. Genetic diversity and population structure of enset (Ensete ventricosum Welw Cheesman) landraces of Gurage zone, Ethiopia. Genetic resources and crop evolution 66(8): 1813–1824.

Getachew S, Mekbib F, Admassu B, Kelemu S, Kidane S, Negisho K, Djikeng A, Nzuki I. 2014. A Look into Genetic Diversity of Enset (Ensete ventricosum (Welw.) Cheesman) Using Transferable Microsatellite Sequences of Banana in Ethiopia. Journal of Crop Improvement 28(2): 159–183.

Glaubitz JC, Casstevens TM, Lu F, Harriman J, Elshire RJ, Sun Q, Buckler ES. 2014. TASSEL-GBS: a high capacity genotyping by sequencing analysis pipeline. PLoS One 9(2): e90346.

Gunesekera B, Torabinejad J, Robinson J, Gillaspy GE. 2007. Inositol polyphosphate 5-phosphatases 1 and 2 are required for regulating seedling growth. Plant Physiology 143(3): 1408–1417.

Guzzon F, Müller JV. 2016. Current availability of seed material of enset (Ensete ventricosum, Musaceae) and its Sub-Saharan wild relatives. Genetic resources and crop evolution 63(2): 185–191.

Harrison J, Moore KA, Paszkiewicz K, Jones T, Grant MR, Ambacheew D, Muzemil S, Studholme DJ. 2014. A Draft Genome Sequence for Ensete ventricosum, the Drought-Tolerant “Tree Against Hunger”. Agronomy 4(1): 13–33.

Heslop-Harrison J, Borrell J, Biswas M, Goodwin M, Blomme G, Schwarzacher T, Wendawek A, Berhanu A, Kallow S, Janssens S. 2019. Enset in Ethiopia: a poorly characterised but resilient starch staple.

Hildebrand E 2001a. Morphological characterization of domestic vs. forest-growing Ensete ventricosum (Welw.) Cheesman, Musaceae, in Sheko district, Bench-Maji Zone, southwest Ethiopia. In: Friis IaR, O ed. Biodiversity Research in the Horn of Africa Region: Royal Danish Academy of Sciences and Letters 287–309.

Hildebrand E 2001b. Morphological characterization of domestic vs. forest-growing Ensete ventricosum (Welw.) Cheesman, Musaceae, in Sheko district, Bench-Maji Zone, southwest Ethiopia. Biodiversity Research in the Horn of Africa Region: Proceedings of the Third International Symposium on the Flora of Ethiopia and Eritrea at the Carlsberg Academy, Copenhagen, August 25–27, 1999: Kgl. Danske Videnskabernes Selskab. 287.

Ishida K, Yamashino T, Mizuno T. 2008. Expression of the cytokinin-induced type-A response regulator gene ARR9 is regulated by the circadian clock in Arabidopsis thaliana. Bioscience, biotechnology, and biochemistry: 0810071089–0810071089.

Jan A, Maruyama K, Todaka D, Kidokoro S, Abo M, Yoshimura E, Shinozaki K, Nakashima K, Yamaguchi-Shinozaki K. 2013. OsTZF1, a CCCH-tandem zinc finger protein, confers delayed senescence and stress tolerance in rice by regulating stress-related genes. Plant Physiology 161(3): 1202–1216.

Jombart T, Devillard S, Balloux F. 2010. Discriminant analysis of principal components: a new method for the analysis of genetically structured populations. BMC genetics 11(1): 94.

Kamanda I, Blay E, Asante I, Danquah A, Ifie B, Parkes E, Kulakow P, Rabbi I, Conteh A, Kamara J. 2020. Genetic diversity of provitamin-A cassava (Manihot esculenta Crantz) in Sierra Leone. Genetic resources and crop evolution: 1–16.

Kang J, Cui H, Jia S, Liu W, Yu R, Wu Z, Wang Z. 2020. Arabidopsis thaliana MLK3, a Plant-specific Casein Kinase 1, Negatively Regulates Flowering and Phosphorylates Histone H3 in Vitro. Genes 11(3): 345.

Konate M, Wilkinson M, Mayne B, Pederson S, Scott E, Berger B, Rodriguez Lopez C. 2018. Salt stress induces non-CG methylation in coding regions of barley seedlings (Hordeum vulgare). Epigenomes 2(2): 12.

Krzywinski MI, Schein JE, Birol I, Connors J, Gascoyne R, Horsman D, Jones SJ, Marra MA. 2009. Circos: an information aesthetic for comparative genomics. Genome research.

Lam H-M, Xu X, Liu X, Chen W, Yang G, Wong F-L, Li M-W, He W, Qin N, Wang B. 2010. Resequencing of 31 wild and cultivated soybean genomes identifies patterns of genetic diversity and selection. Nature genetics 42(12): 1053.

Li N, Li Y. 2014. Ubiquitin-mediated control of seed size in plants. Frontiers in plant science 5: 332.

Li Q-T, Lu X, Song Q-X, Chen H-W, Wei W, Tao J-J, Bian X-H, Shen M, Ma B, Zhang W-K. 2017. Selection for a zinc-finger protein contributes to seed oil increase during soybean domestication. Plant Physiology 173(4): 2208–2224.

Liu K, Muse SV. 2005. PowerMarker: an integrated analysis environment for genetic marker analysis. Bioinformatics 21(9): 2128–2129.

López CMR, Morán P, Lago F, Espiñeira M, Beckmann M, Consuegra S. 2012. Detection and quantification of tissue of origin in salmon and veal products using methylation sensitive AFLPs. Food chemistry 131(4): 1493–1498.

Lotterhos KE, Whitlock MC. 2014. Evaluation of demographic history and neutral parameterization on the performance of FST outlier tests. Molecular Ecology 23(9): 2178–2192.

Lu F, Lipka AE, Glaubitz J, Elshire R, Cherney JH, Casler MD, Buckler ES, Costich DE. 2013. Switchgrass genomic diversity, ploidy, and evolution: novel insights from a network-based SNP discovery protocol. PLoS genetics 9(1).

Ma R, Yuan H, An J, Hao X, Li H. 2018. A Gossypium hirsutum GDSL lipase/hydrolase gene (GhGLIP) appears to be involved in promoting seed growth in Arabidopsis. PLoS One 13(4).

Mandel J, Dechaine J, Marek L, Burke J. 2011. Genetic diversity and population structure in cultivated sunflower and a comparison to its wild progenitor, Helianthus annuus L. Theoretical and Applied Genetics 123(5): 693–704.

Mantel N. 1967. The detection of disease clustering and a generalized regression approach. Cancer research 27(2 Part 1): 209–220.

Martínez-Ainsworth NE, Tenaillon MI. 2016. Superheroes and masterminds of plant domestication. Comptes rendus biologies 339(7): 268–273.

McKey D, Elias M, Pujol B, Duputié A. 2010. The evolutionary ecology of clonally propagated domesticated plants. New Phytologist 186(2): 318–332.

Miller AJ, Schaal BA. 2006. Domestication and the distribution of genetic variation in wild and cultivated populations of the Mesoamerican fruit tree Spondias purpurea L.(Anacardiaceae). Molecular Ecology 15(6): 1467–1480.

Negash A, Tsegaye A, van Treuren R, Visser B. 2002. AFLP analysis of enset clonal diversity in south and southwestern Ethiopia for conservation. Crop Science 42(4): 1105–1111.

Olango TM, Tesfaye B, Catellani M, Pè ME. 2014. Indigenous knowledge, use and on-farm management of enset (Ensete ventricosum (Welw.) Cheesman) diversity in Wolaita, southern Ethiopia. Journal of ethnobiology and ethnomedicine 10(1): 41.

Olango TM, Tesfaye B, Pagnotta MA, Pè ME, Catellani M. 2015. Development of SSR markers and genetic diversity analysis in enset (Ensete ventricosum (Welw.) Cheesman), an orphan food security crop from Southern Ethiopia. BMC genetics 16(1): 98.

Parks DH, Porter M, Churcher S, Wang S, Blouin C, Whalley J, Brooks S, Beiko R. 2009. GenGIS: A geospatial information system for genomic data. Genome research: gr.095612.095109.

Peakall R, Smouse PE. 2012. GenAlEx 6.5: genetic analysis in Excel. Population genetic software for teaching and research—an update. Bioinformatics 28(19): 2537–2539.

Pfeifer B, Wittelsbürger U, Ramos-Onsins SE, Lercher MJ. 2014. PopGenome: an efficient Swiss army knife for population genomic analyses in R. Molecular biology and evolution 31(7): 1929–1936.

Pritchard JK, Stephens M, Donnelly P. 2000. Inference of population structure using multilocus genotype data. Genetics 155(2): 945–959.

Quinlan MB, Quinlan RJ, Dira S. 2014. Sidama Agro-Pastoralism and Ethnobiological Classification of its Primary Plant, Enset (Ensete ventricosum). Ethnobiology Letters 5: 116–125.

Rosenberg NA, Burke T, Elo K, Feldman MW, Freidlin PJ, Groenen MA, Hillel J, Mäki-Tanila A, Tixier-Boichard M, Vignal A. 2001. Empirical evaluation of genetic clustering methods using multilocus genotypes from 20 chicken breeds. Genetics 159(2): 699–713.

Shigeta M. 1990. Folk In-Situ Conservation of Ensete [Ensete ventricosum (Welw.) EE Cheesman]: Toward the Interpretation of Indigenous Agricultural Science of the Ari, Sowthwestern Ethiopia. African Study Monographs 10 (3).

Shumbulo A, Gecho Y, Tora M. 2012. Diversity, Challenges and Potentials of Enset (Ensete ventricosum) Production: In Case of Offa Woreda, Wolaita Zone, Southern Ethiopia. Food Science and Quality Management 7.

Silvertown J. 2008. The evolutionary maintenance of sexual reproduction: evidence from the ecological distribution of asexual reproduction in clonal plants. International journal of plant sciences 169(1): 157–168.

Swain SM, Singh DP, Helliwell CA, Poole AT. 2005. Plants with increased expression of ent-kaurene oxidase are resistant to chemical inhibitors of this gibberellin biosynthesis enzyme. Plant and Cell Physiology 46(2): 284–291.

Tobiaw DC, Bekele E. 2013. Analysis of genetic diversity among cultivated enset (Ensete ventricosum) populations from Essera and Kefficho, southwestern part of Ethiopia using inter simple sequence repeats (ISSRs) marker. African Journal of Biotechnology 10(70): 15697–15709.

Torkamaneh D, Laroche J, Belzile F. 2016. Genome-wide SNP calling from genotyping by sequencing (GBS) data: a comparison of seven pipelines and two sequencing technologies. PLoS One 11(8).

Tsegaye A, Struik P. 2002. Analysis of enset (Ensete ventricosum) indigenous production methods and farm-based biodiversity in major enset-growing regions of southern Ethiopia. Experimental Agriculture 38(03): 291–315.

Uehara TN, Mizutani Y, Kuwata K, Hirota T, Sato A, Mizoi J, Takao S, Matsuo H, Suzuki T, Ito S. 2019. Casein kinase 1 family regulates PRR5 and TOC1 in the Arabidopsis circadian clock. Proceedings of the National Academy of Sciences 116(23): 11528–11536.

Vos P, Hogers R, Bleeker M, Reijans M, Lee Tvd, Hornes M, Friters A, Pot J, Paleman J, Kuiper M. 1995. AFLP: a new technique for DNA fingerprinting. Nucleic acids research 23(21): 4407–4414.

Wen R, Torres-Acosta JA, Pastushok L, Lai X, Pelzer L, Wang H, Xiao W. 2008. Arabidopsis UEV1D promotes lysine-63–linked polyubiquitination and is involved in DNA damage response. The Plant Cell 20(1): 213–227.

Xie H, Konate M, Sai N, Tesfamicael KG, Cavagnaro T, Gilliham M, Breen J, Metcalfe A, Stephen JR, De Bei R. 2017. Global DNA methylation patterns can play a role in defining terroir in grapevine (Vitis vinifera cv. Shiraz). Frontiers in plant science 8: 1860.

Yang H, Wei C-L, Liu H-W, Wu J-L, Li Z-G, Zhang L, Jian J-B, Li Y-Y, Tai Y-L, Zhang J. 2016. Genetic divergence between Camellia sinensis and its wild relatives revealed via genome-wide SNPs from RAD sequencing. PLoS One 11(3): e0151424.

Yemata G. 2020. Ensete ventricosum: A Multipurpose Crop against Hunger in Ethiopia. The Scientific World Journal 2020.

Yemataw Z, Mohamed H, Diro M, Addis T, Blomme G. 2014. Ethnic-based diversity and distribution of enset (Ensete ventricosum) clones in southern Ethiopia.

Zhao D-w, Yang J-b, Yang S-x, Kato K, Luo J-p. 2014. Genetic diversity and domestication origin of tea plant Camellia taliensis (Theaceae) as revealed by microsatellite markers. BMC plant biology 14(1): 14.

Zhong R, Ye Z-H. 2004. Molecular and biochemical characterization of three WD-repeat-domain-containing inositol polyphosphate 5-phosphatases in Arabidopsis thaliana. Plant and Cell Physiology 45(11): 1720–1728.

