## Supplemental Table 4 for "Accumulation of mutations in genes associated with sexual reproduction contributed to the domestication of a vegetatively propagated staple crop, enset"

| **Oligo name** | **Function** | **Sequence** |
| --- | --- | --- |
| *Msp*I adaptor | Reverse Adaptor | CGCTCAGGACTCAT |
| *Msp*I adaptor | Forward Adaptor | GACGATGAGTCCTGAG |
| *Eco*RI adaptor | Reverse Adaptor | AATTGGTACGCAGTCTAC |
| *Eco*RI adaptor | Forward Adaptor | CTCGTAGACTGCGTACC |
| Pre- EcoRI | Pre-selective primer | GACTGCGTACCAATTC**A** |
| Pre- *Msp*I | Pre-selective primer | GATGAGTCCTGAGCGG**C** |
| *Eco*RI Selective Primer | Selective primer | GACTGCGTACCAATTC**ACG** |
| *Msp*I Selective Primer | Selective primer | GATGAGTCCTGAGCGG**CAA** |
| *Msp*I GBS barcoded adaptor | Reverse Adaptor | CGXXXXAGATCGGAAGAGCGTCGTGTAGGGAAAGAGTGT |
| *Msp*I GBS barcoded adaptor | Forward Adaptor | ACACTCTTTCCCTACACGACGCTCTTCCGATCTXXXXX |
| *Eco*RI GBS Y adaptor | Reverse Adaptor | CGAGATCGGAAGAGCGGTTCAGCAGGAATGCCGAG |
| *Eco*RI GBS Y adaptor | Forward Adaptor | CTCGGCATTCCTGCTGAACCGCTCTTCCGATCT |
| *Msp*I GBS primer | Sequencing library primers | AATGATACGGCGACCACCGAGATCTACACTCTTTCCCTACACGACGCTCTTCCGATCT |
| *Eco*RI GBS primer |  | CAAGCAGAAGACGGCATACGAGATCGGTCTCGGCATTCCTGCTGAACCGCTCTTCCGATCT |

**Supplementary Table 5:** Sequences of oligonucleotide used for AFLP. Selective bases in the primers used during the pre-selective and selective amplifications are highlighted in bold. Unique GBS barcode bases are represented as X.
